# Nanorate sequencing reveals the *Arabidopsis* somatic mutation landscape

**DOI:** 10.1101/2025.06.15.659769

**Authors:** Cullan A. Meyer, Brad Nelms, Robert J. Schmitz

**Affiliations:** Department of Genetics, University of Georgia, Athens, Georgia, 30602; Department of Plant Biology, University of Georgia, Athens, Georgia, 30602; The Plant Center, University of Georgia, Athens, Georgia, 30602

**Author notes:** Brad Nelms **Email:**, Robert J. Schmitz **Email:**.

## Abstract

The rate and spectrum of somatic mutations can diverge from that of germline mutations. This is because somatic tissues experience different mutagenic processes than germline tissues. Here, we use nanorate sequencing (NanoSeq) to identify somatic mutations in *Arabidopsis* shoots with high sensitivity. We report a somatic mutation rate of 3.6x10^-8^ mutations/bp, ∼4-5x the germline mutation rate. Somatic mutations displayed elevated signatures consistent with oxidative damage, UV damage, and transcription-coupled nucleotide excision repair. Both somatic and germline mutations were enriched in transposable elements and depleted in genes, but this depletion was greater in germline mutations. Somatic mutation rate correlated with proximity to the centromere, DNA methylation, chromatin accessibility, and gene/TE content, properties which were also largely true of germline mutations. We note DNA methylation and chromatin accessibility have different predicted effects on mutation rate for genic and non-genic regions; DNA methylation associates with a greater increase in mutation rate when in non-genic regions, and accessible chromatin associates with a lower mutation rate in non-genic regions but a higher mutation rate in genic regions. Together, these results characterize key differences and similarities in the genomic distribution of somatic and germline mutations.

## Introduction

The rate of spontaneous mutations can vary between tissue type [1–6], genomic location [7–14], and environmental conditions [15–17]. A better understanding of how often and where new mutations occur would inform many open questions in plant biology and agriculture. For instance, in animals, the somatic mutation rate can be 4-25 times higher than the germline rate [1, 4, 18]; to what degree is this also true in plants? Do plants propagated vegetatively or through tissue culture have a different genomic distribution of mutations compared to plants grown from seed? How might long-lived plants, such as trees, cope with ongoing mutations throughout development?

Answering such questions requires methods to accurately measure the rate and distribution of somatic mutations. This is challenging because current sequencing error rates are ∼10^-3^ [19], orders of magnitude higher than the rate of the spontaneous mutation [1]. One way to address this problem is through deep whole-genome sequencing [2, 3, 12, 13, 20–25]: by focusing on mutations observed multiple times in independent reads, it is possible to distinguish true mutations from sequencing errors. However, the need to repeatedly observe a mutation within a sample limits the analysis to mutations that are relatively abundant. This can bias against detection of mutations acquired later in development which will be rare within a plant. An alternative method without these restrictions is Duplex Sequencing, which can confidently identify mutations from a single DNA molecule [26].

Duplex Sequencing and its successor, nanorate sequencing (NanoSeq), are designed to detect mutations in individual DNA molecules by repeatedly sequencing the top and bottom strand of DNA [1, 27, 28]. True mutations will be present in all PCR duplicates derived from the top strand and the bottom strand. In contrast, PCR errors, sequencing errors, and single-stranded DNA damage are unlikely to be present in duplicates of both strands and can be filtered out. This results in an estimated error rate of 2x10^-7^ errors/bp [1]. NanoSeq improved Duplex Sequencing further by minimizing errors introduced during DNA end repair, which were rare but contributed significantly to error rates [1]. NanoSeq has been used to estimate mutation burden in animal tissues reporting values consistent with single-cell derived colonies, placing its error rate around 5x10^-9^ [1]. One key advantage of NanoSeq and Duplex Sequencing lies in their ability to detect mutations on a molecule-by-molecule basis, allowing for identification of mutations independent of their frequency in the tissue [1, 26]. Duplex Sequencing has shown promise in identifying somatic mutations in *Arabidopsis* [29], but its initial use has returned few mutations in wild type plants (∼1 mutation/plant), limiting the statistical power to ask many important questions. NanoSeq may offer a solution to this recall problem.

Here, we performed NanoSeq on wild-type *Arabidopsis* grown in standard conditions and under UV treatment, identifying 4,155 somatic mutations. We found that 93% of these mutations were unique to a single DNA molecule, supporting an allele frequency below ∼1/200. We compared the untreated plant somatic mutations to germline mutations from mutation accumulation (MA) lines [10] and identified a somatic mutation rate 4-5x that of the germline rate. These somatic mutations also displayed greater signatures of oxidative damage, transcription-coupled nucleotide excision repair, and a UV-like process relative to germline mutations. We then investigated the genome-wide distribution of somatic mutations and found mutations occurred more frequently in transposable elements (TEs), methylated cytosines, and the pericentromere, consistent with observations in MA lines [9, 10]. In addition, we noted accessible chromatin correlated with lower mutation rates outside of genes but higher rates within and upstream of genes. Intriguingly, these patterns were absent in plants treated with UVC, which instead displayed a largely uniform distribution of somatic mutations.

## Results

### Identification of somatic mutations with NanoSeq

We created NanoSeq libraries from the shoots of 8 Col-0 *Arabidopsis* plants (**Fig 1A&B**). Somatic mutations were identified using a custom filtering pipeline (**Fig S1-3 and Methods**). These filters removed single-stranded DNA damage, as well as sequencing, PCR, and alignment errors. We then removed any mutations found in more than one sibling or present in >35% of the reads, since these are likely inherited germline mutations.

**Figure 1:**
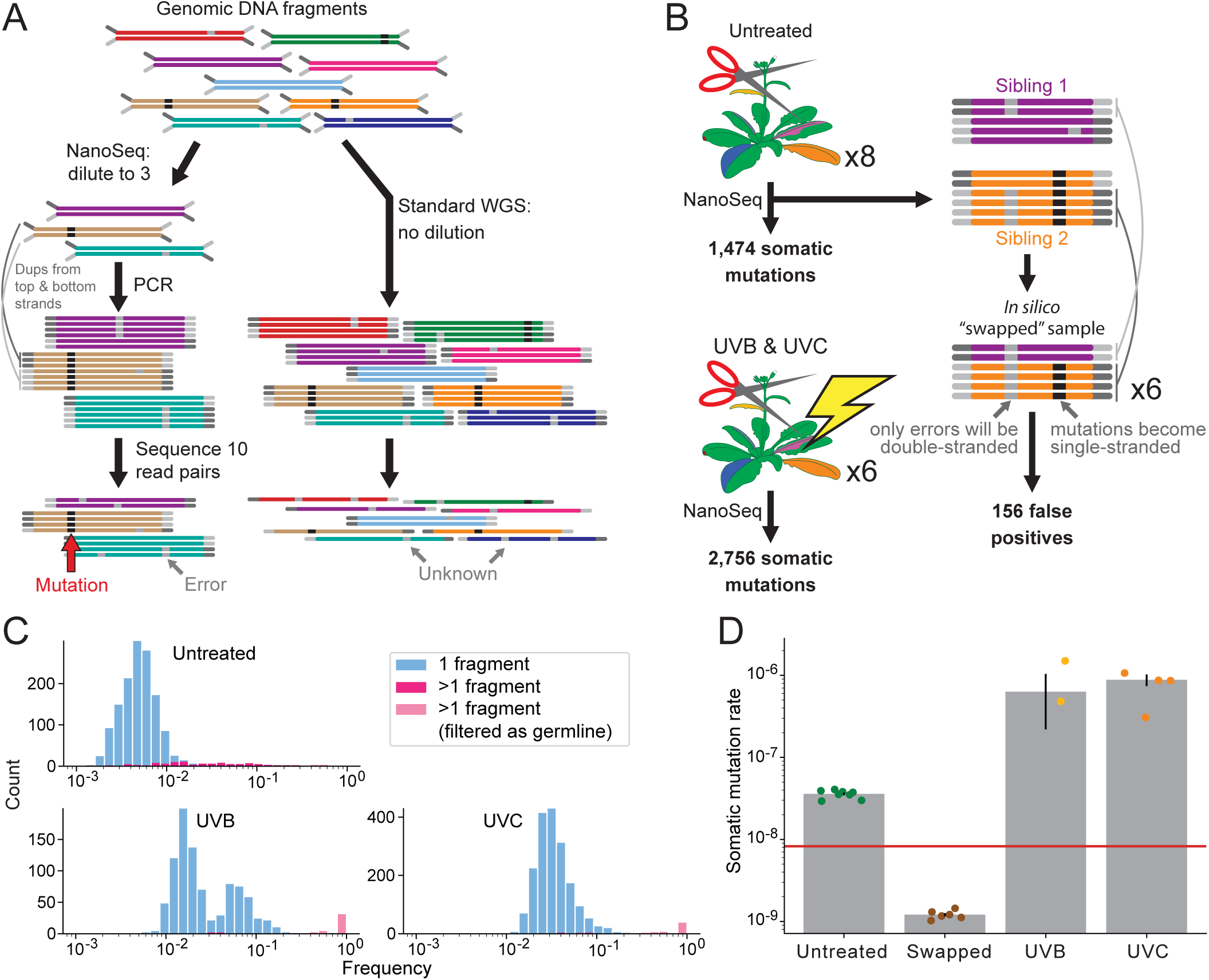
Identification of somatic mutations with NanoSeq. **A**, Schematic of the core difference between NanoSeq and standard whole genome sequencing (WGS) libraries. After fragmentation and adapter ligation of genomic DNA, NanoSeq libraries are diluted to a target number of DNA molecules. PCR and sequencing of the NanoSeq library yield multiple PCR duplicates from each strand of most DNA molecules. Somatic mutations are identified by their presence in all PCR duplicates of a molecule. In contrast, standard WGS has no dilution, so most molecules have only a single read pair, making low abundance mutations indistinguishable from errors. **B**, NanoSeq libraries constructed for this study. Eight untreated and six UV-treated Col-0 sibling *Arabidopsis* were used for NanoSeq libraries and somatic mutations were identified in each. Six *in silico* “swapped” samples were made by combining the top and bottom strand PCR duplicates of molecules from two different untreated NanoSeq libraries. **C**, Frequency of mutations identified by NanoSeq. Mutations found in only a single DNA molecule are colored blue, whereas mutations found in more than one are colored pink. Mutations with a frequency >0.35 were filtered out for further analysis and are colored light pink. **D**, Somatic mutation rates of NanoSeq libraries. Somatic mutation rate was calculated as the mutation count divided by the number of bases in DNA molecules which pass all filters (*i.e.* callable bases). Dots represent replicate plants. The horizontal line is the *Arabidopsis* germline mutation rate [10]. Error bars represent ± one SEM.

We identified 1,425 somatic mutations, including 1,323 single nucleotide variants (SNVs) and 102 small insertions and deletions (indels). Ninety-three percent of mutations were found in only one DNA molecule (**Fig 1C**) but could still be confidently identified as mutations based on duplicate reads from each strand of the DNA duplex. The sites of these single molecule mutations were covered by ∼200 reads, giving a rough upper bound for their frequency of ∼1/200 molecules, or ∼1/100 diploid cells within the plant.

With NanoSeq, each base of a molecule which passes all filters (*i.e.* a callable base) represents an opportunity to detect a mutation. Thus, a sample’s somatic mutation rate was calculated as mutations divided by callable bases (**Methods**). This normalization method reported consistent mutation rates after downsampling the data (**Fig S4**), showing that comparisons between samples of different sequencing depths are possible.

To estimate an error rate for NanoSeq, we generated *in silico* “swapped” libraries, where each molecule consists of a mix of reads from two NanoSeq libraries. This eliminates true mutations, so any identified mutations must be false positives. This method estimated an error rate of 1.2x10^-9^, consistent with the error rate reported for NanoSeq in mammals [1] and 30-fold less than the somatic mutation rate measured in *Arabidopsis* shoots (**Fig. 1D**). To further confirm that we can measure the somatic mutation rate in *Arabidopsis* beyond background errors, we tested whether growing plants under UV would increase the measured mutation rate. Plants were grown with supplemental UVB (N=2 plants) or UVC light (N=4 plants) and subjected to NanoSeq identically as the untreated plants. We identified 2,730 additional somatic mutations in the UV treated plants, with an estimated 18- and 25-fold increase in the measured mutation rate for UVB and UVC (respectively).

The measured somatic mutation rate of the untreated plants was 3.6x10^-8^ mutations/bp. This number represents the average fraction of mutated bases across all DNA in the plants. Multiplying it by the diploid genome size tells us the average diploid somatic cell contains 8.6 mutations which were not present in the zygote. In *Arabidopsis*, the germline mutation rate (SNVs and indels) has been estimated as 7.4-8.25x10^-9^ mutations/bp, or 1.8-1.96 mutations per generation [9, 10]. Thus, the average somatic mutation rate in whole Arabidopsis shoots is ∼4-5x greater than the germline mutation rate.

### SNV spectra of somatic and germline mutations

We next looked for signatures of unique mutagenic processes in somatic mutations compared to germline mutations identified in *Arabidopsis* MA lines [10]. To do this, we visualized the somatic and germline mutation spectra as the rate of insertions, deletions, and each type of SNV (*e.g.* A>G rate = A>G mutations / callable A:T base pairs). Rates were normalized by the genome-wide average mutation rate to make spectra comparable between samples (**Fig 2A**, **Methods**). We then further broke down the SNVs by their 3bp sequence context, normalizing by the frequency of each context in the genome to make rates comparable between contexts (**Fig 2B**).

**Figure 2:**
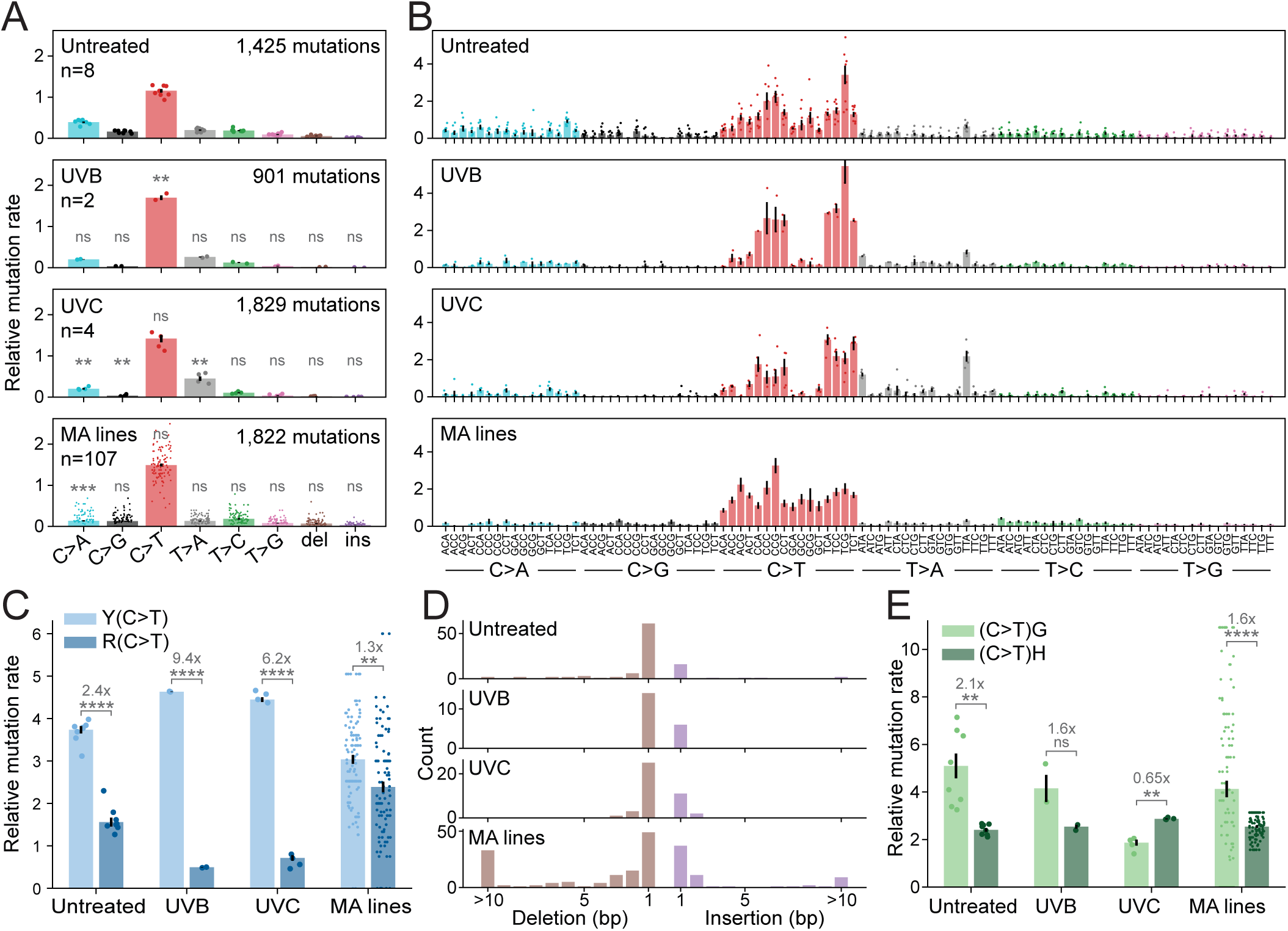
SNV spectra of somatic and germline mutations. **A**, Mutation rate of SNVs and indels by type. The rate for each SNV/indel is the fraction of mutations falling into each category divided by the fraction of callable bases where such a mutation could occur. Significance labels indicate whether the value is significantly different from the same SNV/indel in the untreated samples **B**, SNV mutation rate by 3bp context. Rates are calculated the same as in A. **C**, Comparison of C>T mutation rates in YC vs RC sequence contexts (Y=C/T, R=A/G). **D**, Number of insertions and deletions detected by length. **E**, Comparison of C>T mutation rates in CG vs CH sequence contexts (H=A/C/T). **A-E**, Error bars represent ± one SEM. Dots represent replicate plants/lines. All significance values are from Holm adjusted two-tailed t-tests. *=p<0.05, **=p<0.01, ***=p<0.001, ****=p<0.0001. Untreated=somatic mutations in untreated plants, UVB=somatic mutations in UVB treated plants, UVC=somatic mutations in UVC treated plants, MA lines=germline mutations from published mutation accumulation lines [10].

In the UV-treated samples, C>T SNVs occurred at the highest rate, with YC sites having a far higher rate than RC sites (Y=C/T, R=A/G) (**Fig 2C**). This is consistent with known mechanisms of UV mutagenesis, where the cytosine in UV-induced pyrimidine dimers is spontaneously deaminated to uracil [30]. Prior studies of UV-induced mutations in *Arabidopsis*, yeast, and human cancer display a similar pattern [31–33]. The UV-treated samples also possessed a higher rate of T>A mutations in TTA and ATA contexts. This is likely due to thymidylyl-(3′-5′)- deoxyadenosine, a minor UV-induced photoproduct, and is a signature observed in *Arabidopsis* and yeast, but not humans [32, 34, 35].

When comparing somatic mutations from the untreated plants to the germline mutations, we noted a difference in the rate of C>A mutations. Of the germline SNVs, 5.6% were C>A, but this increased to 15.3% in the somatic mutations (p<0.001, Holm adjusted two-tailed t-test). In addition, the C>T mutation rate in the untreated samples was 2.4x higher in YC contexts than RC contexts, significantly greater than the 1.3x difference in the germline mutations (**Fig 2C**, p<0.01, Holm adjusted one-sample two-tailed t-test). Given that the plants were grown indoors under fluorescent lights, we expect UV levels to be very low, so the YC>YT somatic mutations may represent a mutational process other than UV. While fewer large indels (>10bp) were observed in the somatic mutations (**Fig 2D**), our mapping pipeline for NanoSeq is not optimized for large indels and likely under reports these. In most other regards, the somatic and germline mutation spectra were comparable, being dominated by C>T mutations and displaying higher rates of C>T mutation at CG sites (**Fig 2E**).

### Global distribution of somatic and germline mutations

We next aimed to characterize the distribution of somatic mutations within the genome compared to germline mutations. In addition to the MA line dataset, we included two polymorphism datasets; the first was 13,792,559 polymorphisms from wild accessions of the 1001 Genomes Project [36]. Second, we used NanoSeq to identify 624,111 fixed polymorphisms in Ler-0—an accession within the 1001 Genomes dataset. These polymorphism datasets provide much greater power than the MA lines but represent historical mutations observed in a population rather than selfing lines. For simplicity, we will refer to the somatic mutations, MA line mutations, and polymorphisms collectively as “mutations”.

Mutation rate was calculated in sliding 2 megabase (Mb) windows across the genome (**Fig 3A**). Here, mutation rate refers to the number of mutations within the window divided by the callable bases within the window (or the number of callable sites for the MA line, Ler-0, and 1001 Genomes datasets, **Methods**). Thus, the mutation rate factors in the mappability and sequencing coverage of the window. For most datasets, mutation rate was highest near the centromere and lower on the chromosome arms. To quantify this relationship, we plotted the mutation rate of non-overlapping 1Mb bins against their distance to the centromere.

**Figure 3:**
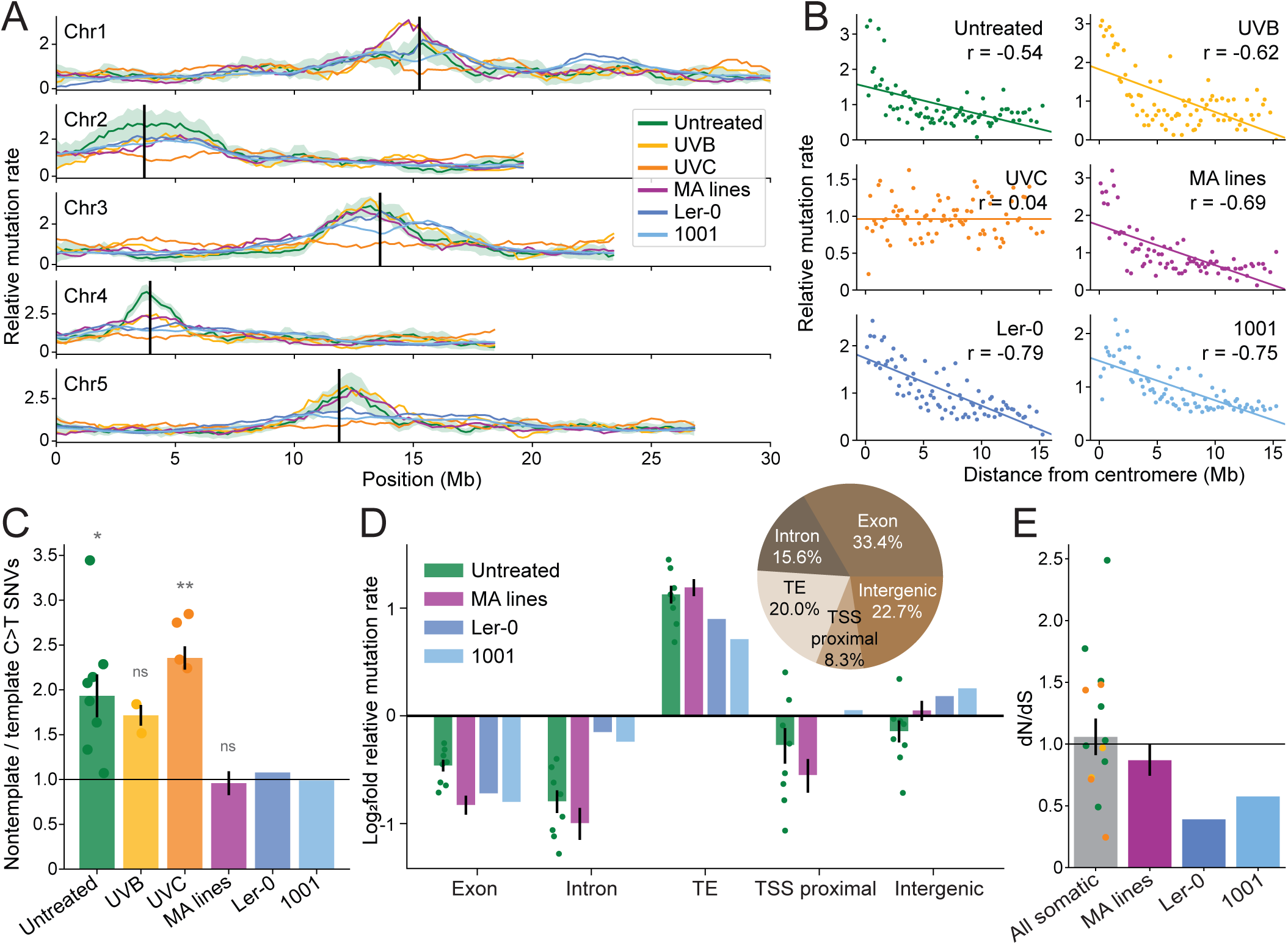
Global distribution of somatic and germline mutations. **A**, Mutation rate relative to the genome average across each chromosome. Mutation rate is calculated for sliding 2Mb windows as mutations divided by callable base pairs (untreated, UVB, and UVC) or callable sites (MA lines, Ler-0, and 1001). A value of 1 indicates the genome-wide average mutation rate for the sample. Vertical black lines indicate the centromere positions [39]. Shaded area around the untreated line represents a 95% confidence interval from bootstrapping samples. **B**, Correlation between distance to centromere and mutation rate. Each dot represents one non-overlapping 1Mb window. Linear regression line (ordinary least squares) and correlation coefficients are displayed. **C**, Ratio of nontemplate:template strand C>T mutations. Significance labels indicate whether the mean is significantly greater than one (Holm adjusted one-sample one-tailed t-test) **D**, Log_2_ fold relative mutation rate by genomic region. A value of 0 indicates the region has the same mutation rate as the genome-wide average. Pie chart indicates the proportion of the *Arabidopsis* genome classified as each region. **E**, Mutation rate in nonsynonymous sites relative to synonymous sites in each dataset. **C-E**, Error bars represent ± one SEM. For panels C & E, SEM of the MA lines was calculated by bootstrapping lines. Dots represent replicate plants. *=p<0.05, **=p<0.01. Untreated=somatic mutations in untreated Col-0, UVB=somatic mutations in UVB treated Col-0, UVC=somatic mutations in UVC treated Col-0, MA lines= germline mutations from published Col-0 mutation accumulation lines [10], Ler-0=polymorphisms in Ler-0 identified with NanoSeq, 1001= polymorphisms from published dataset of 1,135 wild accessions [36].

For most datasets, we observed a negative correlation between mutation rate and distance to centromere (**Fig 3B**); the one exception was the UVC-treated samples, where the mutation rate was more uniform across the chromosomes. We hypothesized that the uniform mutation rate in UVC, but not UVB, samples could be due to differential activity of transcription-coupled nucleotide excision repair (TC-NER). This pathway is known to repair UV damage only on the template strands of transcribed genes and so should be more active in the chromosome arms [37]. To test for signatures of TC-NER activity, we compared rates of C>T mutations when the cytosine was on the template vs nontemplate strand (**Fig 3C**). Contrary to our hypothesis, both UVB and UVC samples showed a strong signature of TC-NER activity, with C>T mutations roughly twice as common on the nontemplate strand. We also noted signatures of TC-NER activity in the untreated somatic mutations but not in the other datasets. Thus, TC-NER activity appears to be influencing the somatic mutation rate but not the germline mutation rate, and the uniform mutation rate in UVC samples is not a product of diminished TC-NER activity.

The pericentromere has more TEs and fewer genes than the chromosome arms. Thus, a higher mutation rate in the pericentromere could reflect a higher mutation rate in TEs. We categorized every site in the genome as one of five regions—exon, intron, TE, transcription start site (TSS) proximal, or intergenic. TE regions were defined using a published TE annotation [38], TSS proximal as 1 to 500bp upstream all transcription start sites, and intergenic as anything not falling into the other four regions. For each region, the mutation rate was calculated relative to the genome-wide average rate (**Fig 3D**). In most datasets, exons and introns were depleted for mutations, whereas TEs were enriched, with TEs possessing 3.2x the somatic mutation rate of genes. However, genes were more depleted for MA line mutations than for somatic mutations, as seen by a lower relative mutation rate (p<0.05, Holm adjusted two-tailed Welch’s t-test). Only the UVC treated samples did not have a significantly lower mutation rate in genes (p=0.49, one-sample t-test), instead displaying a largely uniform mutation rate across the five regions (**Fig S7**). In addition, TSS proximal regions had a lower mutation rate than the intergenic regions in the MA lines (p<0.01, Holm adjusted two-tailed t-test) and polymorphism datasets, but this difference was not significant in the somatic mutations (p=0.53, two-tailed t-test). Lastly, the Ler- 0 and 1001 datasets displayed a substantially higher mutation rate in introns compared to exons, which may be due to the influence of selection on these datasets.

To look for signatures of purifying selection, we calculated a global dN/dS value for each dataset as the ratio of mutation rate in nonsynonymous to synonymous (4-fold degenerate) sites. As expected, the Ler-0 and 1001 genomes datasets had a dN/dS significantly lower than one, but the somatic mutations and MA line mutations did not (**Fig 3E**). This suggests purifying selection is too weak to detect in both the somatic and MA line mutations and is unlikely to be driving their observed distribution.

### Somatic mutation rate correlates with centromere proximity, DNA methylation, and chromatin accessibility

Previous work in *Arabidopsis* has found that the germline mutation rate correlates with proximity to the centromere and DNA methylation [10]. However, it is not known whether these factors influence somatic and germline mutation rates to the same degree nor whether they correlate with somatic mutation rate at all.

To test whether somatic mutation rate correlates with proximity to the centromere independent of TE content, we compared mutation rates of pericentromeric genes and TEs (<5Mb from the centromere) to distal genes and TEs (>5Mb from the centromere) (**Fig 4A**). In all four datasets, pericentromeric genes had a higher mutation rate than their distal counterparts, but this difference was not significant for the somatic mutations nor the MA lines (p=0.08, p=0.36, Holm adjusted two-tailed t-test). Pericentromeric TEs had 1.9x the somatic mutation rate of distal TEs (p<0.01, Holm adjusted two-tailed t-test), and this was significantly greater than the 1.1x difference in MA lines (p<0.05, Holm adjusted one-sample two-tailed t-test). Thus, proximity to the centromere associated with a greater increase in TE mutation rates in somatic contexts than in germline contexts.

**Figure 4:**
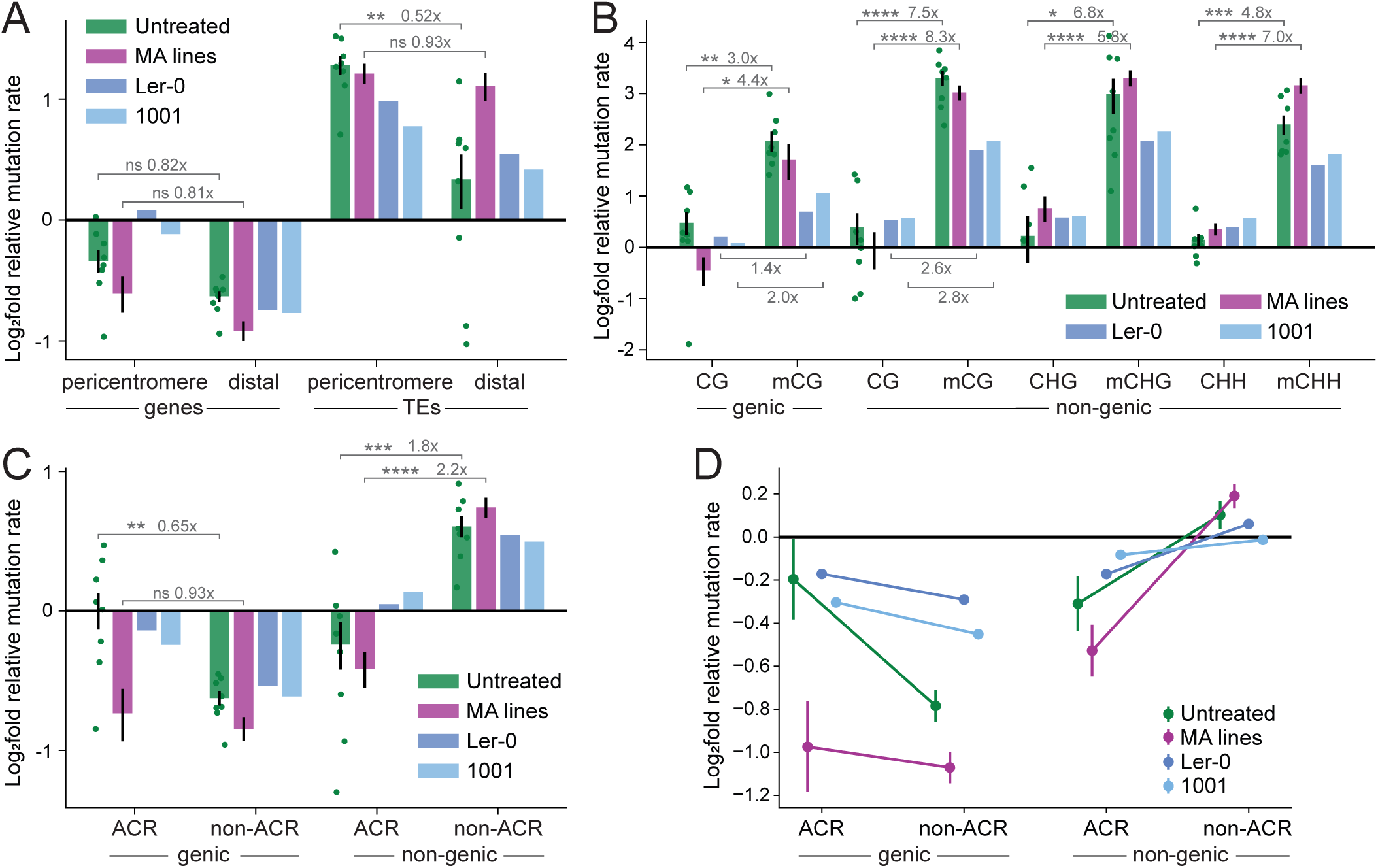
Somatic mutation rate correlates with centromere proximity, DNA methylation, and chromatin accessibility. **A**, Log_2_ fold relative mutation rate of genes and TEs in pericentromeric (<5Mb from the centromere) vs distal (>5Mb) regions. **B**, Log_2_ fold relative mutation rate of genic CG sites and non-genic CG, CHG, and CHH sites by methylation status in Col-0. **C**, Log_2_ fold relative mutation rate of ACR overlapping vs non-overlapping regions. Two regions are considered, genic and non-genic. Each bar represents the mutation rate in the intersection of the specified region and ACR or non-ACR space. **A-C**, Error bars represent ± one SEM. Dots represent replicate plants. Significance labels are for Holm adjusted two-tailed t-tests. *=p<0.05, **=p<0.01, ***=p<0.001, ****=p<0.0001. **D**, Linear/logistic regression predictions of mutation rate in ACRs vs non-ACRs. Points represent the predicted mutation rate of the model under fixed C:G content, DNA methylation, distance to centromere, and nonsynonymous site content. Error bars represent ± one SEM calculated using the delta method. Untreated=somatic mutations in untreated Col-0, MA lines= germline mutations from published Col-0 mutation accumulation lines [10], Ler-0=polymorphisms in Ler-0 identified with NanoSeq, 1001= polymorphisms from published dataset of 1,135 wild accessions [36].

In *Arabidopsis*, DNA methylation is primarily found in TEs and genes. TEs have high levels of cytosine methylation in all sequence contexts—CG, CHG, and CHH (where H=A/C/T) [40, 41]. In contrast, most genes are unmethylated (∼80%) or methylated only in CG contexts (∼17%), though a small fraction are methylated in all sequenced contexts (∼3%) [42]. To investigate the correlation of DNA methylation with mutation rate, we used published bisulfite sequencing data to identify methylated cytosines in Col-0 [43]. We compared methylated and unmethylated genic CG sites and found the methylated sites had a significantly higher mutation rate in all four datasets (**Fig 4B**). The same was true for non-genic CG, CHG, and CHH sites. However, in all four datasets, methylation in non-genic regions associated with a greater increase in mutation rate than in genic regions. We saw little difference between somatic and MA line mutations, suggesting DNA methylation has a similar mutagenic effect in somatic and germline contexts.

In human cancers, chromatin accessibility has been shown to anticorrelate with mutation rate [44]. To test whether this is also the case in *Arabidopsis*, we made use of published ATAC-seq data to identify accessible chromatin regions (ACRs) in Col-0 [45]. Looking only within non-genic regions, we calculated mutation rate of ARCs vs non-ACRs (**Fig 4C**). In all four datasets, non-genic regions had a lower mutation rate if they overlapped an ACR, consistent with the observation in human cancers [44]. We then made the same comparison but looked only within genic regions. Intriguingly, we now observed the opposite pattern, where genic regions overlapping an ACR had a higher mutation rate than other genic regions. This difference was only significant for the somatic mutations and not the MA line mutations (p<0.01 & p=0.60, Holm adjusted two-tailed t-test). Thus, accessible chromatin is associated with a lower somatic mutation rate outside of genes, but a higher rate within genes.

ACRs are generally unmethylated and less common near the centromere [45–48], so we wondered whether these factors were driving the correlation of non-genic ACRs with a lower mutation rate. To test this, we made linear/logistic regression models which identify independent associations of various factors with mutation rate. For each dataset, we fit a model of mutation rate per genomic site which considered whether the site was a C:G pair, methylated, accessible, in the pericentromere, in a gene, and—for the Ler-0 and 1001 datasets only—whether it was nonsynonymous (**Methods**). Lastly, we considered a potential interaction term between chromatin accessibility and genic regions. All four models predicted a higher mutation rate for pericentromeric and methylated sites, consistent with previous analyses (**Table S1**). In addition, they predict ACRs to have a lower mutation rate outside of genes and a higher mutation rate within genes, in agreement with Figure 4C (**Fig 4D, Table S1**).

## Discussion

### The *Arabidopsis* somatic mutation rate is 4-5x the germline mutation rate

We used NanoSeq to identify somatic mutations in untreated and UV-treated *Arabidopsis* independent of their frequency within the plant. An average of 178 mutations/plant were identified, representing a major improvement in recall over a previous Duplex Sequencing study [29]. 93% of these somatic mutations were observed in only one DNA molecule, placing their frequency at somewhere less than 1/100 cells. We report a somatic mutation rate 4-5x higher than the germline mutation rate, in other words, the average *Arabidopsis* shoot cell contains 4-5x as many new mutations as a zygote of the next generation. This difference is less than what is observed for most human somatic tissues, which accumulate mutations 4-25x as rapidly as the paternal germline [1, 4].

The higher somatic mutation rate could be explained if the average shoot cell has undergone more cell divisions since the zygote than the sperm or egg. DNA replication is known to be mutagenic, as it introduces polymerase errors and propagates DNA damage into double-stranded mutations [49]. However, measurements of telomere shortening in *Arabidopsis* telomerase mutants estimate that rosette leaves undergo fewer cell divisions than the progeny, not more [50]. Another possibility would be that the cell lineage leading to the next generation has a lower average mutation rate per mitotic division. In humans and mice, per cell division mutation rate estimates are an order of magnitude higher in fibroblasts than the germline [51]. Even nondividing neurons and smooth muscle have higher mutation rates than the mitotically active male germline [1]. Thus, the higher somatic mutation rate in *Arabidopsis* may not be the result of more cell divisions but instead an increase in mutagenic processes and/or less efficient DNA repair.

### Somatic mutations possess signatures of oxidative damage and a UV-like process

We found that C>A and YC>YT mutations were more common in somatic than germline contexts. As the somatic mutations likely include those accumulated outside the meristem, these signatures may represent mutagenic processes more active in terminal tissues. One potential source of the C>A mutations is reactive oxygen species, which react with DNA to form 8-oxo-7,8-dihydroguanine (8-oxoG) and other lesions [52]. 8-oxoG then mispairs with adenine, producing C>A mutations [52]. Reactive oxygen species are produced by chloroplast metabolism, resulting in much higher levels of ROS in leaves than roots [53–55]. Thus, the increased rate of C>A somatic mutations may be a consequence of greater photosynthetic activity in the leaves compared to the meristem. The origin of the YC>YT somatic mutations is much less clear. This mutation signature is often associated with UV-induced pyrimidine dimers, but the untreated plants were not exposed to sunlight [30]. This signature could represent some other mutagenic process or unexpected UV emission from the fluorescent grow lights.

We also noted a strong signal of TC-NER in the C>T somatic mutations, whereas this signature was absent in the germline mutations. The simple explanation for this is that TC-NER is more active in terminal tissues than the meristem. An analogous situation occurs in *C. elegans*, where TC-NER is more active in somatic tissues, and global genome NER (GG-NER) is more active in the germline [56–58]. Perhaps *Arabidopsis* has adopted a similar strategy, where GG-NER maintains the integrity of the whole genome in the meristem, while terminal tissues only require maintenance of transcribed genes and so use TC-NER. An alternative explanation for the absence of a TC-NER signature in the germline mutations is that the germline C>T mutations are caused by a different type of DNA damage which TC-NER cannot repair.

### Global distribution of somatic mutations

Somatic and germline mutations were enriched near the centromere and in TEs, while genes were depleted. This pattern has previously been noted in *Arabidopsis* as dependent on mismatch repair activity, suggesting mismatch repair is more efficient within genes [12, 59]. We note that the depletion of mutations in genes was slightly greater in germline than somatic contexts. This may be caused by differential mismatch repair activity in the meristem and terminal tissues. Another explanation is purifying selection; MA lines are theorized to experience selection against strong deleterious mutations which may have no consequences in heterozygous somatic contexts [60]. However, we lacked the statistical power to detect any purifying selection in the germline and somatic mutations.

Intriguingly, the UVC treated samples were the only plants to not display an enrichment for mutations in the centromere nor a depletion in genes. The UVB and UVC treatments differed in the wavelength (311nm vs 254nm) and duration (8h/day vs ∼3s/day), with UVC treatments being far more concentrated. We suspect the UVC treatments overwhelmed repair pathways which had time to preferentially repair damage in genes during UVB treatment. UV damage can be repaired by TC-NER, GG-NER, and photoreactivation [37, 61]. Strong signatures of TC-NER activity were observed in the UVC sample, indicating this pathway was not overwhelmed. Studies in yeast have revealed that photoreactivation occurs significantly faster in nucleosome free DNA [62, 63]. Thus, we hypothesize photoreactivation preferentially repaired genes and euchromatin in the UVB samples but was overwhelmed by the UVC treatment. It is also worth noting that for the UVC mutation rate to be uniform while TC-NER is active, UVC must have induced more damage in genes than non-genic regions. This is the case in yeast, where UVC induces more pyrimidine dimers on the template strand of genes than flanking regions [64].

### Proximity to the centromere, DNA methylation, and chromatin accessibility correlate with somatic mutation rate

Methylated cytosines had higher somatic and germline mutation rates, consistent with known properties of ∼2-5x faster deamination rates and ∼2.6x higher rates of DNA polymerase errors at methylated cytosines [9, 10, 65–68]. *Arabidopsis* and most flowering plants have CG methylation in the body of many genes, but this methylation has no known function [69–72]. Thus, it is puzzling why plants would maintain this mutagenic modification in genes [73, 74]. Our data suggest DNA methylation within genes is less mutagenic than methylation in non-genic regions, reducing the fitness cost of maintaining gene body methylation.

We found that pericentromeric TEs had a higher somatic mutation rate than distal TEs, which may be explained by replication timing. In both *Arabidopsis* and animals, heterochromatic domains—like the pericentromere—replicate late in S phase [75, 76]. These late replicating regions have higher mutation rates in human cancers, dependent on functional mismatch repair activity [77, 78]. *Arabidopsis* mismatch repair may also be less functional late in S phase, when the pericentromere is being replicated.

Within non-genic regions, chromatin accessibility correlated with a lower somatic and germline mutation rate, consistent with what is seen in human cancers [44]. In yeast, repair of alkylation and UV damage is faster at sites where the DNA minor groove faces away from the nucleosome [64, 79]. Thus, we suspect non-genic ACRs in *Arabidopsis* have lower mutation rates because they are more accessible to repair factors. In contrast, genic regions had higher somatic mutation rate when overlapping an ACR. As most of these ACRs are present near the site of transcription initiation, their elevated mutation rate may represent transcription-associated mutagenesis, where unwinding of the DNA makes it more susceptible to damage [80].

## Conclusion

We used NanoSeq to generate an unbiased measurement of somatic mutations in *Arabidopsis.* Somatic mutations accumulated at a faster rate than germline mutations and possessed signatures of oxidative damage, TC-NER activity, and a UV-like mutational process. Somatic mutation rate correlated with gene/TE content, proximity to the centromere, DNA methylation, and chromatin accessibility, similar to germline mutations. Deducing the causal effects of these factors on mutation rate will require further studies under genetic perturbation.

## Supplemental Figures

**Supplemental Figure 1:**
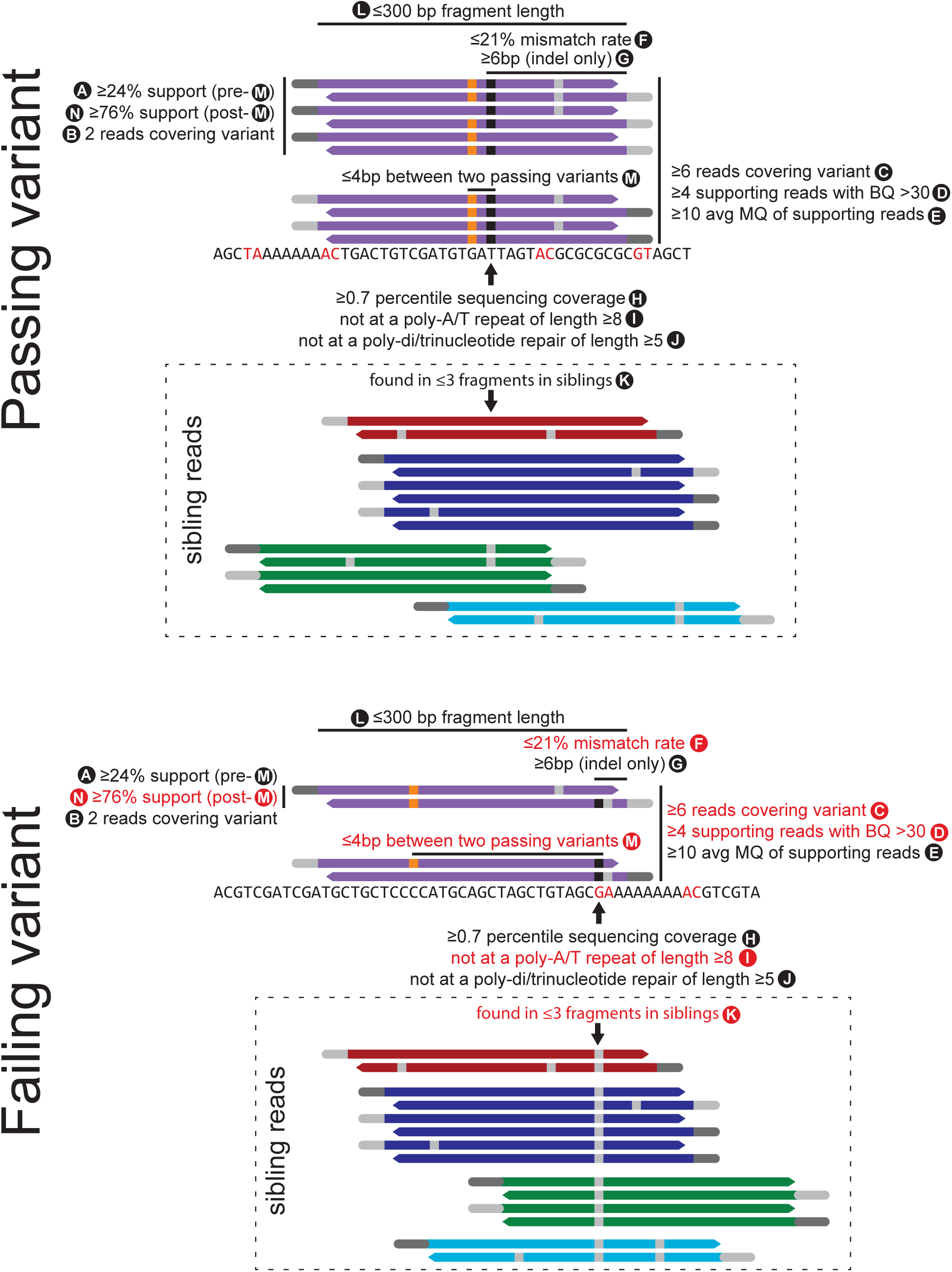
Filters used to call somatic mutations. **Top**, A variant which passes all filter is depicted as a black rectangle within the purple fragment. Each purple line represents a sequencing read. Read pairs are interleaved, with read1s possessing a dark grey adapter and read2s possessing a light grey adapter. The top set of purple lines represent PCR duplicates which used the top strand of the original DNA fragment as a template, whereas the bottom set of purple lines used the bottom strand as a template. Filters are labeled A-M. The genomic sequence is shown below the purple reads with blacklisted sites (filters G-I) colored red. The orange variant also passes all filters. **Bottom**, A variant which fails several filters is depicted in black. The filters which the variant fails are colored red. The orange variant also fails filters. **F**, Mismatch rate refers to the number of SNVs + the length of all indels between the variant and fragment end which pass filters A-E divided by the distance between the variant and fragment end. **G**, Indels must be ≥6bp from each fragment end. **H**, Sequencing coverage of every site in the genome was calculated as the number of reads in all untreated plants which overlap it. **K**, A fragment within a sibling library is considered to contain the variant if it is supported by both ≥2 and ≥79% of reads. **M**, If any two variants within the same fragment pass filters A-L and are >4bp apart, all variants within the fragment are discarded. **N**, This filter is the same as filter “A”, just with a higher threshold and applied only after filter “M”.

**Supplemental Figure 2:**
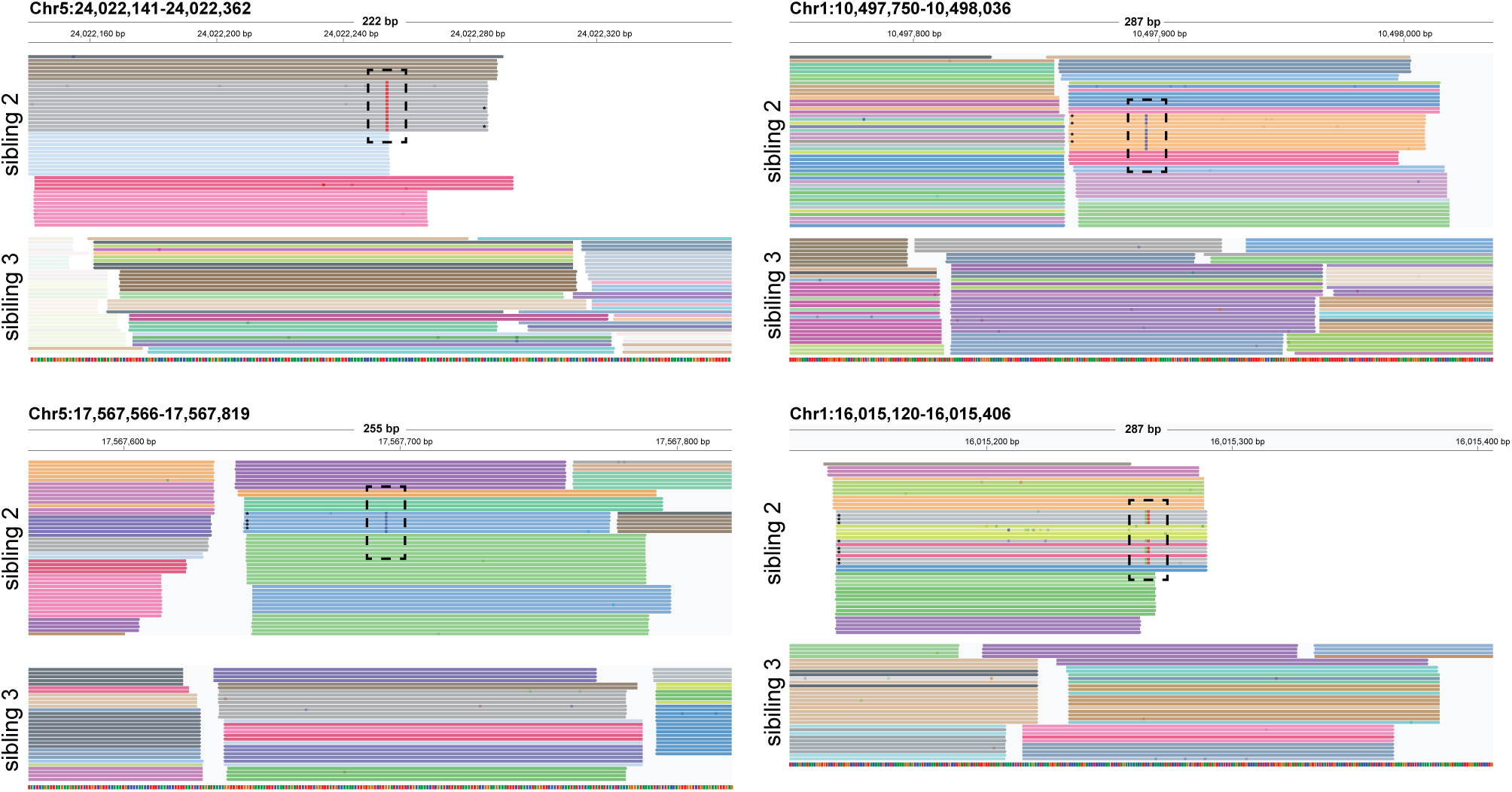
Representative genome browser views of somatic mutations. Four somatic mutations were randomly sampled from those identified in untreated sibling two. Each line represents a sequencing read. Reads which are PCR duplicates possess the same color. Somatic mutations are within dotted outlines. Reads which are PCR duplicates of the bottom DNA strand (read1 is in the reverse direction) are labeled with a star. Reads from sibling three are also shown as a control. Transparent reads and non-reference bases indicate low mapping and base quality respectively.

**Supplemental Figure 3:**
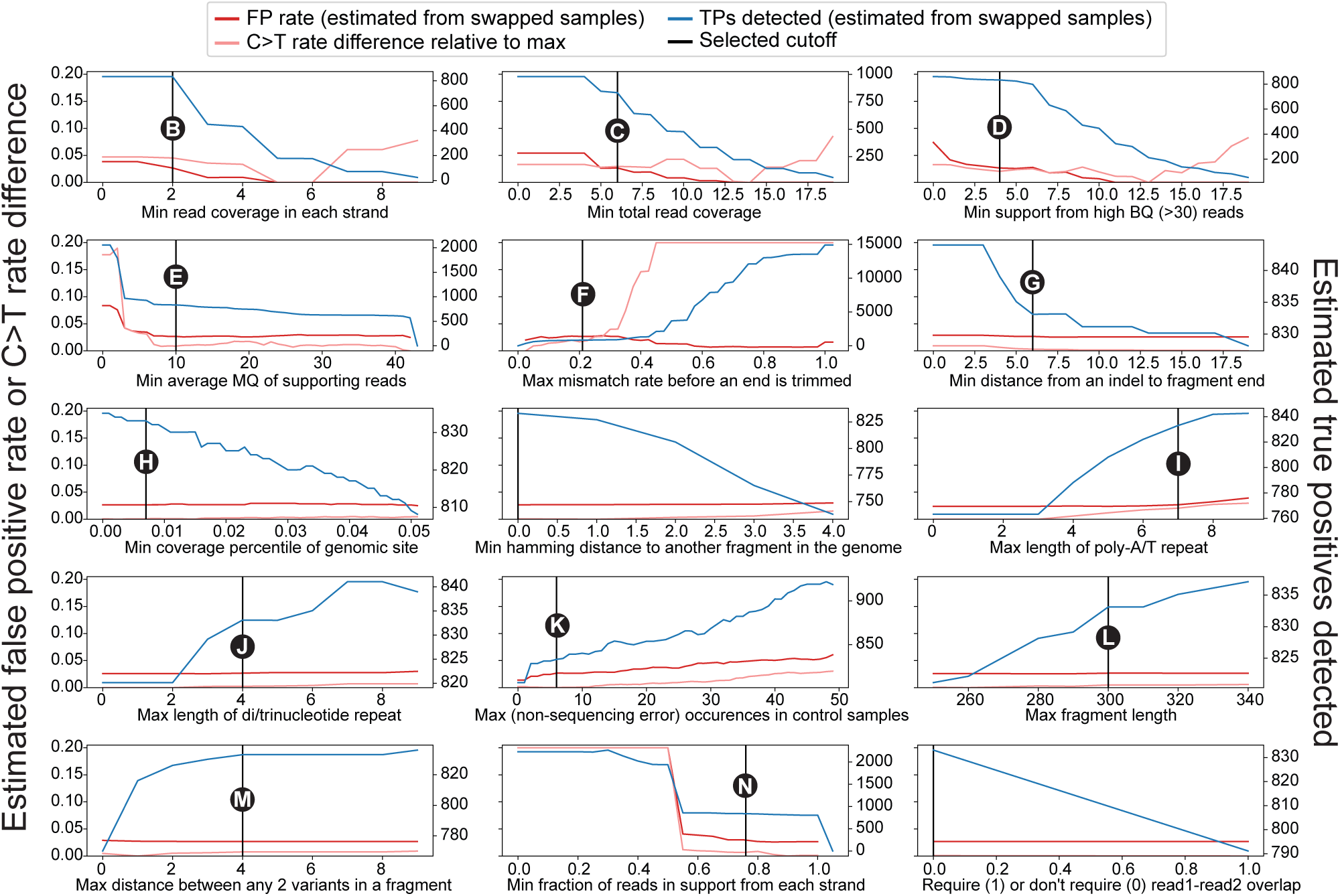
Filter threshold decisions. Unfiltered variants were identified in untreated samples one through four and 3 “swapped” samples. The somatic mutation filters were then applied as described (**Fig S1 and Methods**), except one filter threshold was allowed to vary. Each panel represents varying of one filter threshold. False positive rate was estimated as the somatic mutation rate (number of passing variants / callable coverage) of the swapped samples divided by that of the untreated samples. Number of true positives was estimated as the number of passing variants * (1 – false positive rate). The C>T rate was calculated for each threshold as the fraction of passing variants which were C>T SNVs. The C>T rate difference was the maximum C>T rate within the panel minus the C>T rate at the threshold. X-axis labels indicate which filter threshold is being varied. Black lines indicate the threshold used for all analyses. The black circles link each filter to its depiction in Figure S1. Two filters which were not used in the final analysis have no black circle.

**Supplemental Figure 4:**
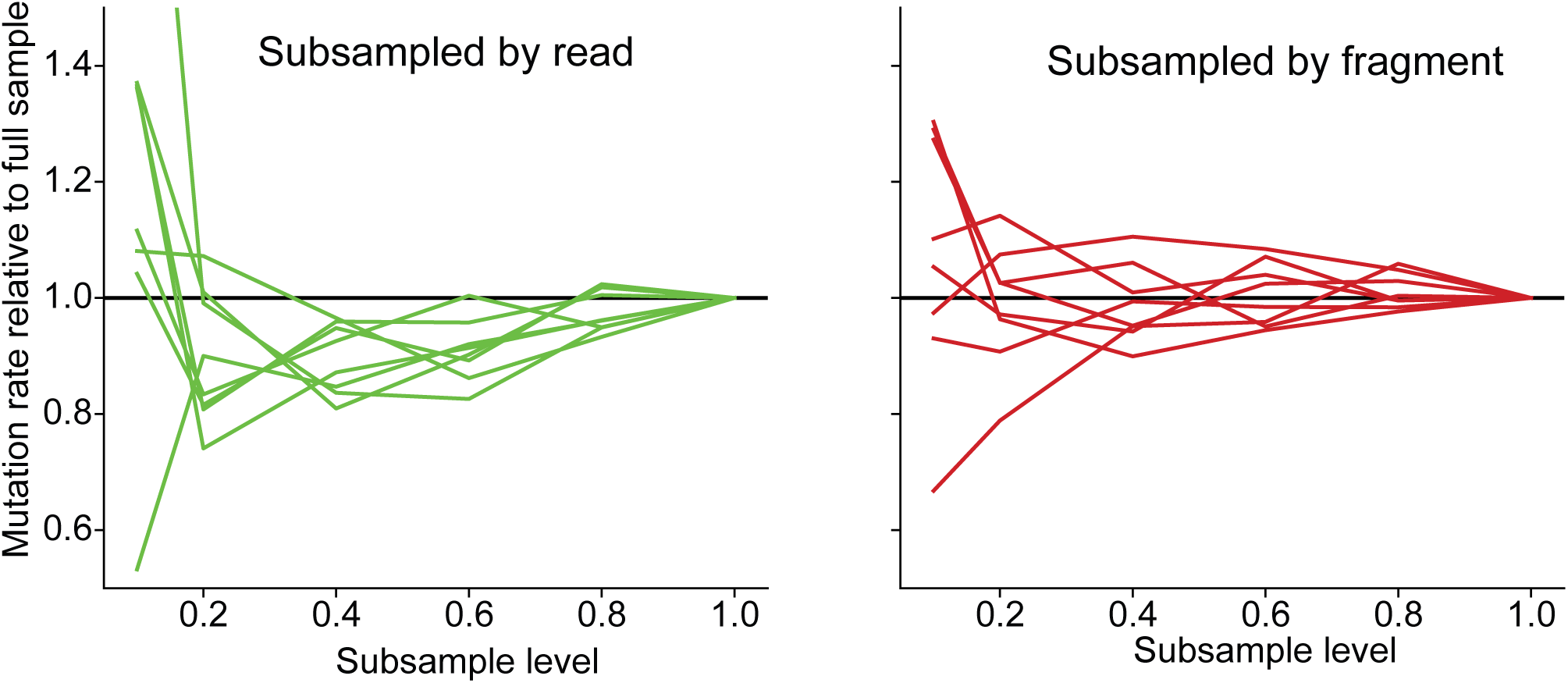
NanoSeq somatic mutation rate is stable across subsampling. For each of the eight untreated NanoSeq libraries, randomly subsampled libraries with 80%, 60%, 40%, 20%, and 10% of the original library read count were created. Callable coverage was recalculated for each subsampled library and used to generate a somatic mutation rate relative to the full sample rate. **Left**, reads were randomly sampled from NanoSeq libraries, with no consideration for fragment. Thus, subsampled libraries have a lower read:fragment ratio, representing under-diluted (under-sequenced) NanoSeq libraries. NanoSeq libraries have a slight trend toward underestimating mutation rate when under-diluted. **Right**, fragments were randomly sampled from NanoSeq libraries, so all selected fragments in the subsampled libraries have the same number of PCR duplicates as in the full libraries. Thus, subsampled libraries have the same read:fragment ratio, representing properly diluted libraries. Properly diluted libraries have no bias toward underestimating nor overestimating somatic mutation rate and remain within 0.8-1.2x the full sample rate at a subsample level of 20%.

**Supplemental Figure 5:**
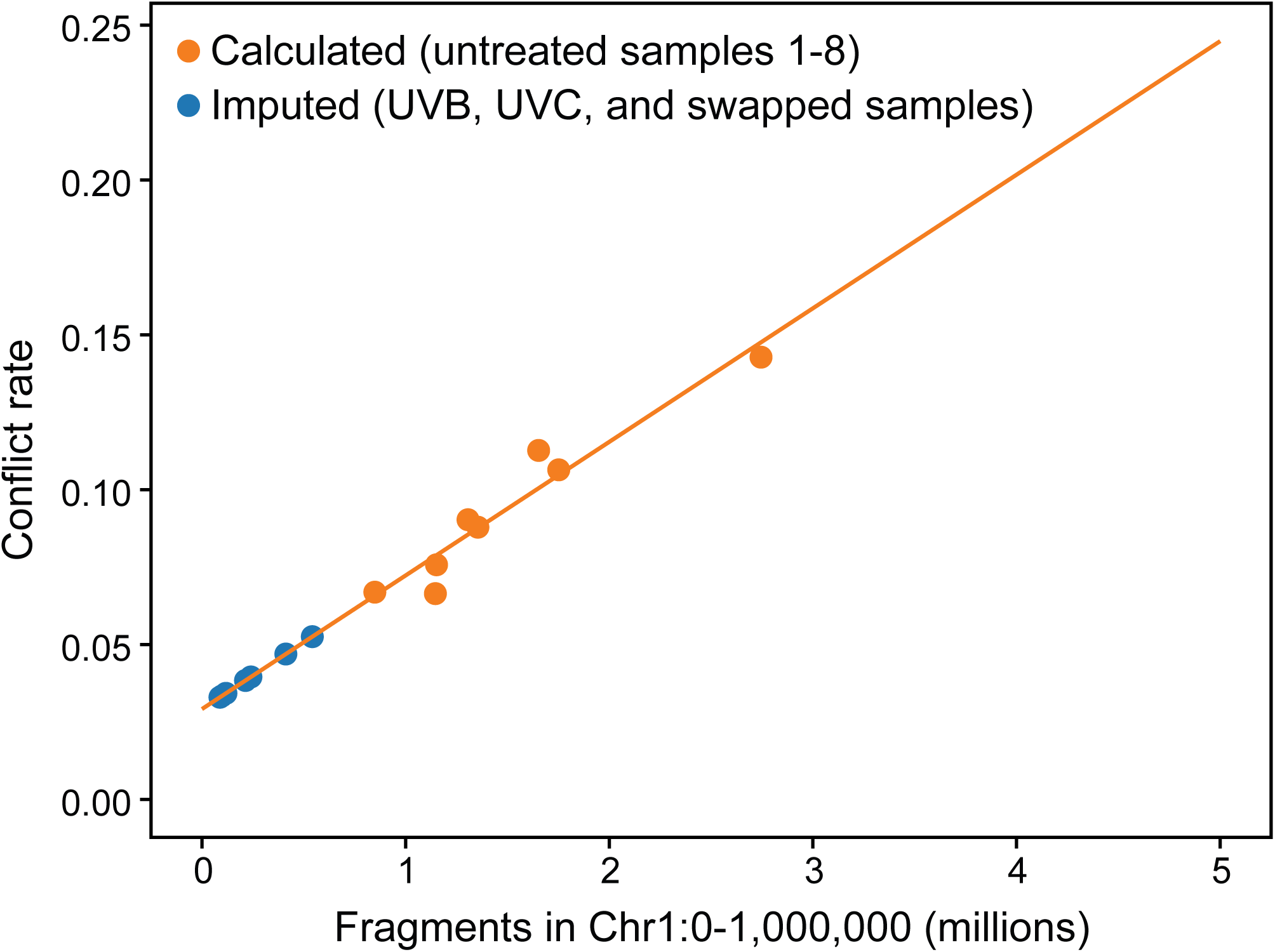
NanoSeq fragment conflict rate scales linearly with fragments sequenced. Fragment conflict rate was calculated for all untreated NanoSeq libraries as described in Methods (orange dots). For samples with no replicates (UVB, UVC, and swapped, blue dots), conflict rate could not be calculated and was instead imputed based on their fragment number and the best fit line of the untreated libraries.

**Supplemental Figure 6:**
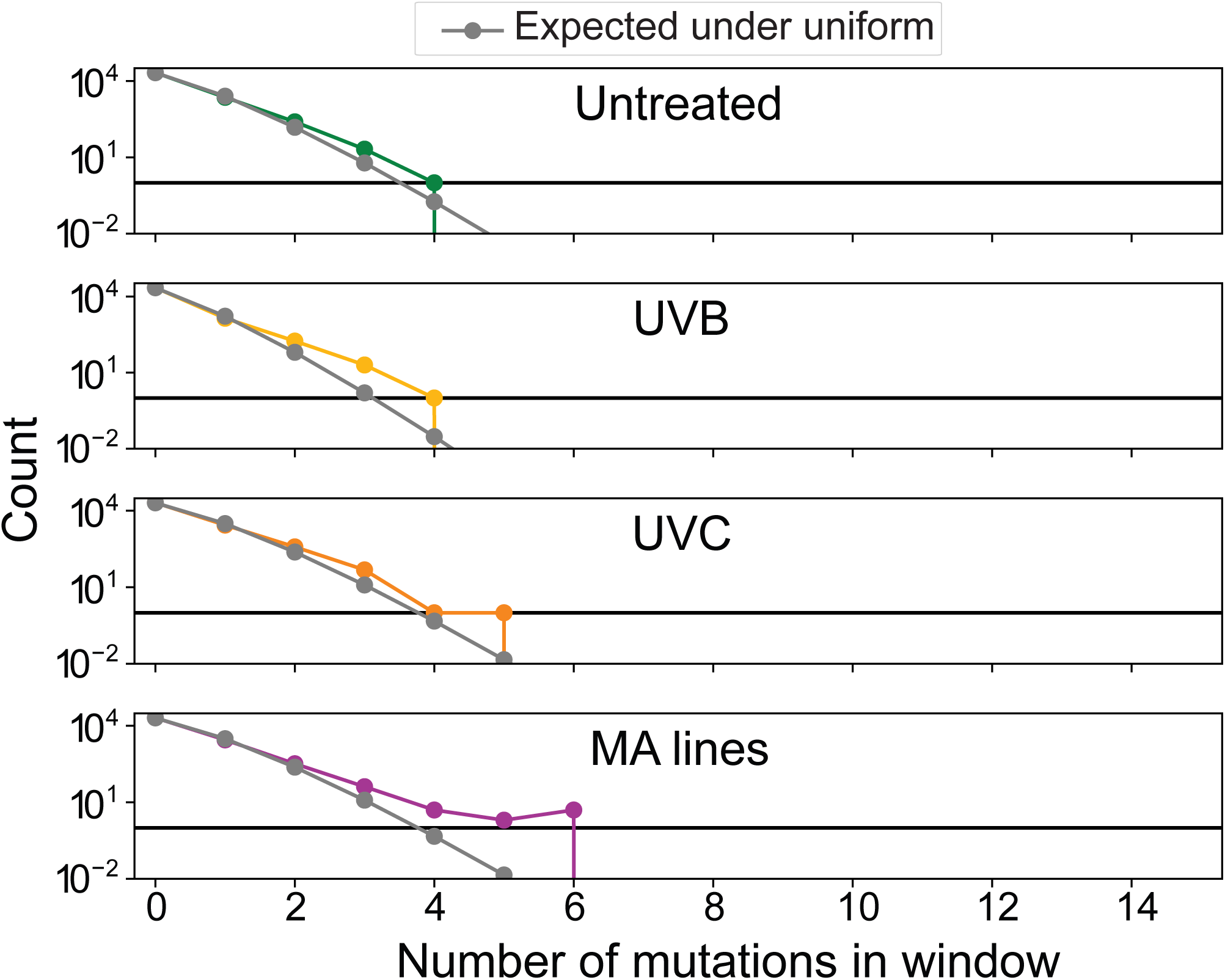
No signs of mutation hotspots in somatic mutations. The genome was segmented into 10kb half-overlapping windows and the number of mutations in each window was counted. Colored lines represent histograms of the number of mutations per window. Grey lines represent the expected number of windows assuming each site in the genome is equally likely to be mutated (mutations per window follows a binomial distribution). Horizontal black line is placed at a value of one. No samples have any windows with more than 6 mutations.

**Supplemental Figure 7:**
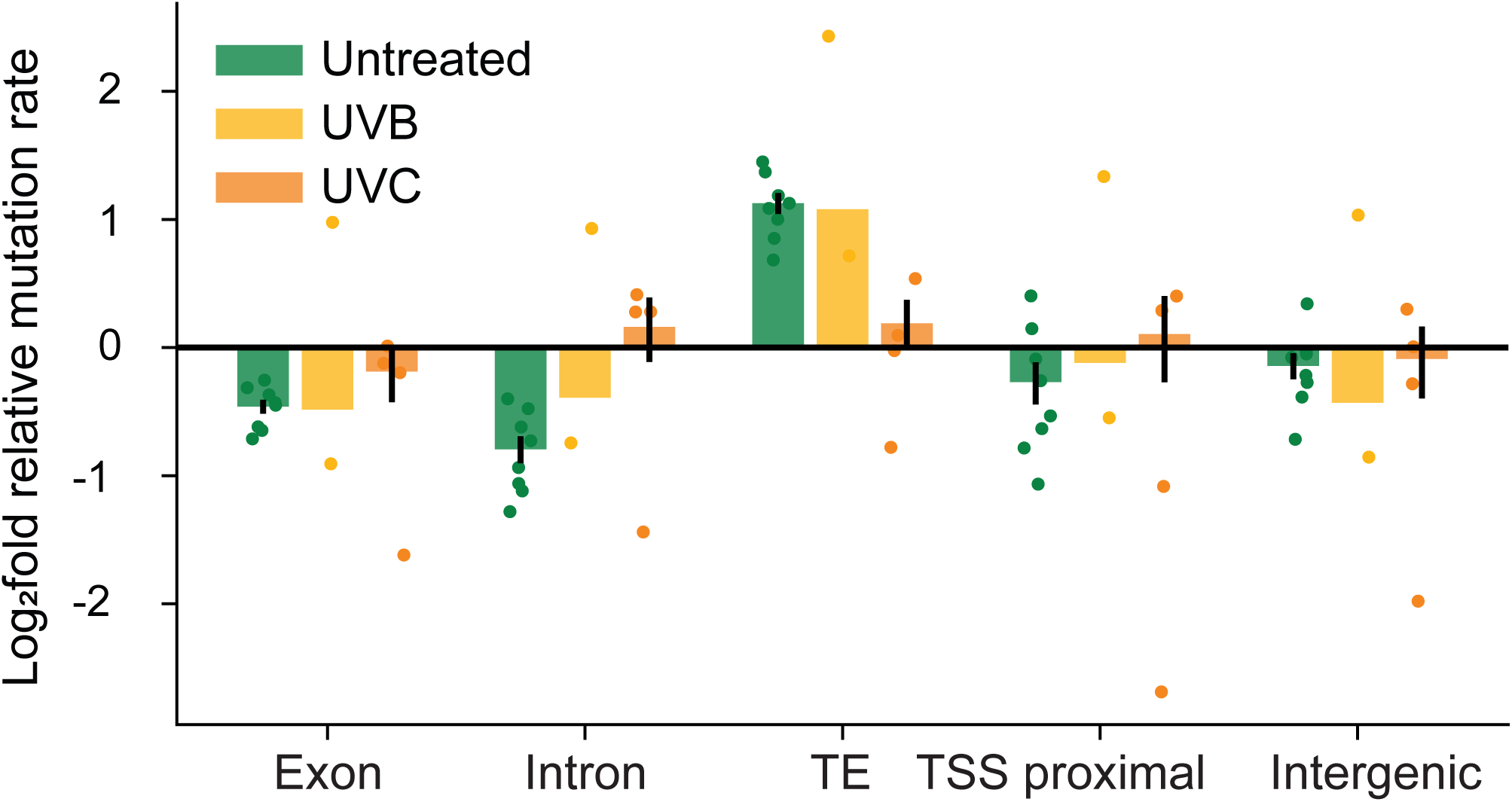
UVC treated samples are not depleted for genic mutations. Equivalent to Figure 3D. Log_2_ fold relative substation rate by genomic region. A value of 0 indicates the region has the same mutation rate as the genome-wide average. Dots represent individual somatic mutation samples. Error bars represent ± one SEM. Untreated=somatic mutations in untreated plants, UVB=somatic mutations in UVB treated plants, UVC=somatic mutations in UVC treated plants.

**Supplemental Figure 8:**
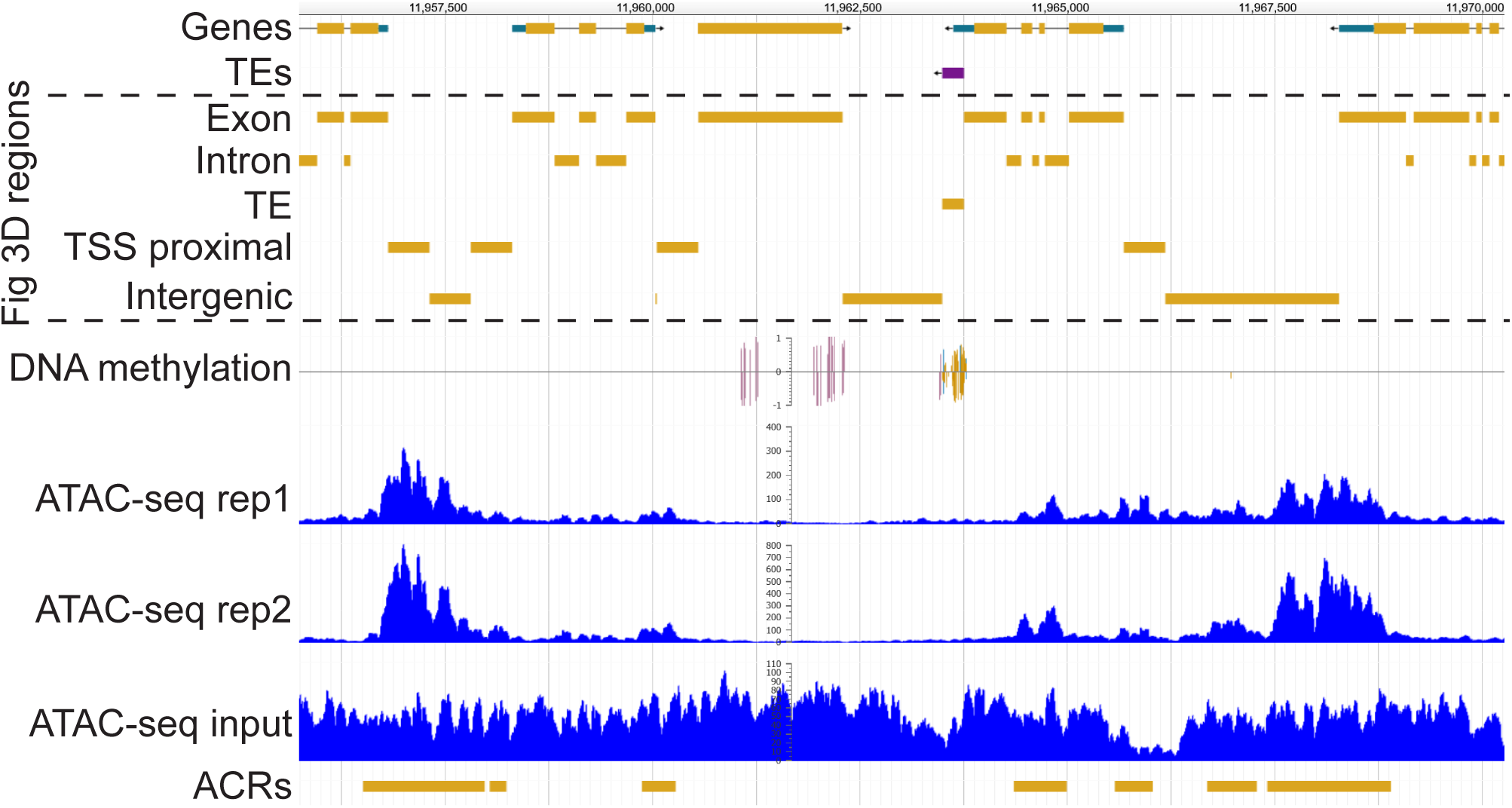
Bisulfite and ATAC-seq genome browser screenshot. Genome browser screenshots of the five genomic regions used in Figure 3D, processed bisulfite sequencing data, and processed ATAC-seq data. **Top to bottom**, gene annotations from TAIR10 [81]; TE annotations from Panda and Slotkin [38]; 5 genomic regions used in Figure 3D; bisulfite sequencing data used for Figure 4B [43], bars indicate methylated cytosines (purple=mCG, blue=mCHG, yellow=mCHH), bar height indicates the fraction of reads supporting methylation; ATAC-seq data used for Figure 4C [45], bigwigs of two replicates and one input sample are shown, as is the final set of accessible chromatin regions used.

## Methods

### Plant material

All plants were grown in Sungro soil containing Osmocote fertilizer at 21°C under 16 hours of light. All Col-0 plants were siblings. Untreated Col-0 and Ler-0 were grown under a mix of Philips F96T8/TL841 PLUS and Sylvania Octron Eco fluorescent lamps. UVC treated Col-0 were grown under the same conditions, but treated with UVC from a Spectroline Select Series UV Crosslinker at multiple points throughout the course of growth. The number and intensity of UVC treatments varied between the four plants (10x0.02J/cm^2^, 10x0.05J/cm^2^, 4x0.1J/cm^2^, 2x0.2J/cm^2^). UVB treated Col-0 were grown under Sylvania Octron Eco and a Philips TL 20W/01Narrowband UVB lamp. The UVB lamp was ∼20cm from the plants and on for every other hour of the 16-hour day.

All plants were harvested at the opening of the first flower. For the Col-0 plants, flowers and flower buds were removed and the remaining shoot was flash frozen. For the Ler-0 plant, a single leaf was flash frozen.

### NanoSeq

Each frozen tissue sample was ground with a mortar and pestle. DNA was isolated with a DNeasy Plant Mini Kit (Qiagen, 69106) following manufacturer’s instructions. DNA was fragmented to 150bp using a Covaris E220 Evolution Focused-ultrasonicator by the Georgia Genomic and Bioinformatics Core. 200ng of fragmented DNA was size selected using in-house AMPure XP magnetic beads (0.79 beads:sample ratio for lower and 2.15 for upper selection). End blunting was performed by incubating in a reaction of 1x S1 nuclease buffer and 10units S1 nuclease (Thermo, EN0321) for 30 min at RT. The reaction was stopped by addition of 3µL EDTA, cleaned up with magnetic beads (1.8 ratio), and eluted in 32µL 10mM TrisHCl (pH 8). A-tailing and nick blocking was performed by first adding 5µL 10x *E. coli* DNA ligase buffer (NEB, M0205S), 4.5µL 10mM ATP, and 1uL 10units/µL T4 Polynucleotide Kinase (NEB, M0201S) and incubating at 37°C for 30 min. Samples were then placed on ice, 1uL 10unit/µL *E. coli* DNA ligase (NEB, M0205S) was added, and samples were incubated at 16°C for 30 min. A mix of 1.75µL NF H_2_O, 0.25µL 10mM dATP, 0.5µL ddCTP, 0.5µL ddGTP, 0.5µL ddTTP (Cytiva, 27204501), and 3µL 5units/µL Klenow fragment (3’ -> 5’ exo-) (NEB, M0212S) was added and incubated at 37°C for 30 min. Samples were cleaned up with magnetic beads (1.8 ratio) and 2µL 8µM iTruSeq adapter stubs (sequence below) were added. Adapter ligation was performed in a 25µL reaction of 1x T4 DNA ligase buffer and 20units/µL T4 DNA ligase (NEB, M0202S). Samples were incubated at 16°C for 16 hours and cleaned up with two rounds of magnetic beads (1.4 ratio).

DNA concentration was measured by Quibit and a portion of each library was diluted to 0.1ng/µL. Three dilutions were used to make a 0.0125ng/µL dilution. These dilutions and three 0.1ng/µL dilutions of a previously sequenced library were run on qPCR in triplicate 10µL reactions of 1x Luna qPCR universal master mix (NEB, M3003S), 1.5µM Illumina i5 indexed primer, and 1.5µM Illumina i7 indexed primer (sequences below). The qPCR program was 95°C (60s), 35x (95°C (15s), 60°C (30s)). Cq values of new and previously sequenced libraries were compared to determine the optimal volume of each sample to use per million read pairs sequenced, such that the number of callable fragments is maximized. An optimal volume of sample for the amount of sequencing planned was used for further steps. For untreated Col-0 samples, samples were split in three to reduce the chance of DNA molecules with the same alignment start and end position being present in a single library.

Samples were amplified in a 50µL reaction of 0.33µM Illumina i5 indexed primer, 0.33µM Illumina i7 indexed primer, 200µM dNTPs, 1x Phusion HF Buffer, and 0.04units/µL Phusion HF DNA Polymerase (NEB, M0530S). PCR program was 98°C (60s), 2x (98°C (15s), 50°C (120s), 72°C (15s)), 11x (98°C (15s), 60°C (30s), 72°C (15s)). Samples were cleaned up with magnetic beads (1.4 ratio) and sequenced 150bp paired end reads on the NovaSeq X series platform 25B flow cell.

iTruSeq stub sequences: ACACTCTTTCCCTACACGACGCTCTTCCGATCT and /5phos/GATCGGAAGAGCACACGTCTGAACTCCAGTCAC

Illumina i5 and i7 indexed primers: AATGATACGGCGACCACCGAGATCTACACNNNNNNNNACACTCTTTCCCTAC and CAAGCAGAAGACGGCATACGAGATNNNNNNNNGTGACTGGAGTTCAG where Ns are the sample index

### Filtering and alignment of reads

Sequencing reads were trimmed and filtered with fastp using default parameters [82]. Reads were aligned to the TAIR10 reference genome using Bowtie2 with -X 800 [81, 83]. Optical duplicates were then marked using SAMtools fixmate -m, Sambamba sort, and SAMtools markdup -d 2500 [84, 85]. Optical duplicates, non-concordantly mapped, and ambiguously mapped reads were filtered using sambamba view --filter mapping_quality ≥ 1 and proper_pair and ([dt] == null or [dt] != ‘SQ’). Replicate libraries generated from the same plant were then merged using SAMtools merge -r to produce the final filtered BAM files. These commands were run as a Snakemake pipeline [86].

### *In silico* generation of a “swapped” control

The filtered BAM files of the untreated plants were used to generate “swapped” libraries using a custom script. For each set of PCR duplicates present in more than one plant (same fragment alignment start and end site), one set of top strand PCR duplicates (read1 on the forward strand) and one set of bottom strand PCR duplicates (read1 on the reverse strand) were selected at random from two different plants. These selected PCR duplicates were output as a single swapped library. The process was then repeated five more times to generate six swapped BAM files.

### Filtering somatic mutations

A set of unfiltered variants were identified using a custom script. These variants were then filtered by passing the following requirements: ≥24% of reads in the top strand duplicates and ≥24% of reads in the bottom strand duplicates support the variant; ≥6 duplicate reads cover the variant position; ≥2 top strand duplicates and ≥2 bottom strand duplicates cover the variant; ≥4 duplicate reads support the variant with a BQ >30 at the variant base (if SNV); ≥10 average MQ of supporting reads; ≤0.21 mismatches/bp between the variant and each fragment end (this removes all variants <5bp from the fragment end); variant is ≥6bp from fragment ends (applied to indels only); variant position is ≥0.7 percentile in total read coverage across all untreated BAMs (*i.e.* removes genomic positions with low read coverage); variant position is not the start/end of a poly-A nor poly-T repeat of length ≥8; variant position is not the start/end of a dinucleotide nor trinucleotide repeat of ≥5 repeating units; variant is supported by the majority of reads in ≤3 sets of PCR duplicates across the eight untreated libraries; variant is in a fragment of length ≤300bp; there are ≤4bp between any 2 variants in the same set of PCR duplicates passing all previous filters (this filter was not applied to the Ler-0 sample); ≥76% of reads in the top strand duplicates and ≥76% of reads in the bottom strand duplicates support the variant (**Fig S1**). Variants present in a sibling plant were filtered as likely germline mutations. We determined the cutoffs for these filters by varying each one and assessing how it affected the number of variants identified in untreated samples and swapped samples as well as the fraction of variants which were C>T (**Fig S3**).

The frequency of each mutation in the sample was calculated as the fraction of duplicate sets covering the variant site which had >76% of reads supporting the variant. For the Ler-0 sample, only variants with a frequency >0.35 were retained. For all other samples, only variants with a frequency <0.35 were retained.

### Calculation of callable coverage and somatic mutation rate

For the untreated and UV treated samples, callable coverage was calculated per site as the number of opportunities a somatic mutation could have been observed and passed all filters. Each set of PCR duplicates was considered “callable” if it had ≥1 bottom strand, ≥1 top strand, and ≥3 total read pairs, as well as average MQ ≥10 and fragment length ≤300bp. Then, each base within a callable fragment contributed to callable coverage if it overlapped ≥2 reads from the bottom strand, ≥2 reads from the top strand, and ≥6 total reads, was ≥5bp from a fragment end, and was not in any blacklisted regions of the genome. Blacklisted regions were those with <0.7 percentile total read coverage across all untreated BAMs, the starts/ends of ≥8bp poly-A and poly-T repeats, and the starts/ends of ≥5 length di/trinucleotide repeats. For the Ler-0 sample, callable coverage was limited to a max of one, so every site had callable coverage of either 0 (uncallable) or 1 (callable).

To calculate the overall somatic mutation rate of a sample, the callable coverage of every site in the nuclear genome was summed and divided by a predicted rate of fragment conflicts. The conflict rate represents the chance each fragment in the library has another fragment with the same alignment start and end site, which may preclude mutation identification in that fragment. Conflict rate was estimated by randomly selecting half of the fragments from two replicate libraries of the same plant and calculating a “half” conflict rate as the fraction of selected fragments which conflict with another selected fragment. The full conflict rate was then calculated as 1 – (1 – half conflict rate)^2^. For samples with no replicate libraries, the conflict rate was imputed using the other samples assuming a linear relationship between conflict rate and fragments in the library (**Fig S8**). Thus, the overall somatic mutation rate was calculated as the number of mutations / (callable coverage * (1 – conflict rate)). Conflict rate was not considered when calculating somatic mutation rate for specific genomic regions nor when calculating mutation spectra (**Fig 2-4**).

### Processing of published mutation/polymorphism datasets

For the MA line dataset, we downloaded mutations from Weng, Becker [10]. These were homozygous mutations present in one of 107 Col-0 lines propagated for 25 generations of single seed descent. For the 1001 genomes dataset, SNVs and short indels in 1,135 wild *Arabidopsis* accessions were retrieved from 1001genomes.org [36]. We discarded mutations in both datasets if they were present at a position with no callable coverage in the untreated first sibling NanoSeq library. As an estimate of callable coverage in these datasets, we took the callable coverage of the untreated first sibling NanoSeq library and limited the coverage per site to one, so every site had callable coverage of either 0 (uncallable) or 1 (callable).

### Calculation of mutation spectra

A mutation rate was calculated for each SNV type, 3bp context, insertions, and deletions as the count of that mutation type divided by the callable coverage of sites which could harbor that mutation (*e.g.* coverage of all C:G pairs for C>T mutations, coverage of all ACA sites for ACA>AAA mutations, and coverage of all sites for indels). This mutation rate was then divided by the genome-wide average mutation rate for that sample to get the relative mutation rate.

### DNA methylation and chromatin accessibility

Bisulfite sequencing data of Col-0 were downloaded from SRR2922654 Bewick, Ji [43]. Sequencing reads were trimmed and filtered with fastp using default parameters [82]. Methylation files were generated using methylpy --trim-reads False --merge-by-max-mapq True --min-mapq 1 --min-qual-score 1 [87]. Methylated sites were called using METHimpute [88]. Cytosines with an “Intermediate” methylation status were not considered methylated nor unmethylated for generating Figure 4B.

Two replicates of Col-0 ATAC-seq data were downloaded from PRJNA527732 [45]. Sequencing reads were trimmed and filtered with fastp using default parameters [82]. Reads were aligned to the TAIR10 reference genome using Bowtie2 with -X 800 [83]. Peaks were called for each replicate using MACS2 callpeak -g 1.1e8 -q 0.7 --nomodel --extsize 200 --shift -100 [89]. The union of peaks in the two replicates was used as the final ACR set.

### Linear/logistic regression models of mutation rate

For each of the four datasets, we constructed a linear/logistic regression model using the Statsmodels package (**Table S1**) [90]. Each genomic site had the following predictor variables with values of 0 (false) or 1 (true), C:G pair, methylated, within an ACR, within 5Mb of the centromere, within a gene, nonsynonymous coding site, and within both an ACR and a gene (ACR * gene). The nonsynonymous variable was only considered in the models for the Ler-0 and 1001 datasets. The response variable was modeled differently for each dataset; for the untreated dataset, a generalized linear model was used, and the response variable was modeled as a binomial distribution where trials=callable coverage and successes=mutations at the site (generalized linear model). For the MA line and Ler-0 datasets, a logistic regression model was used, and the response variable was whether a mutation was present at the site. For the 1001 genomes dataset, a generalized linear model was used, and the response variable was modeled as a Poisson distribution where successes=mutations at the site (this dataset could have >1 mutation per site). Each model was fit using the genomic sites with >0 callable coverage in the dataset. To calculate mutation rate in Figure 4D, we predicted the mutation rate for a site with the ACR, gene, and ACR*gene variables set to 111, 010, 100, or 000 and all other predictors set to their average value across all sites.

## Data Availability

NanoSeq library sequencing data can be found on NCBI SRA database under the BioProject accession PRJNA1247547.

## Acknowledgements

We would like to thank Mark Minow for suggestions on data processing and visualization, Bruce Martin for advice on regression methods, and Yangyang Xu for performing DNA extractions. The Georgia Advanced Computing Resource Center provided the computational resources required for data analysis. This research was supported by the National Science Foundation (MCB-2242696) and the University of Georgia Office of Research to R.J.S. as well as the National Institute for General Medical Sciences of the National Institute of Health to C.A.M. (1T32GM142623) and B.N. (R35GM151237).

